# Speckle rheological spectroscopy reveals wideband viscoelastic spectra of biological tissues

**DOI:** 10.1101/2023.06.08.544037

**Authors:** Nichaluk Leartprapun, Ziqian Zeng, Zeinab Hajjarian, Veerle Bossuyt, Seemantini K. Nadkarni

**Affiliations:** Wellman Center for Photomedicine, Massachusetts General Hospital, Harvard Medical School, Boston, MA 02114 USA; Department of Pathology, Massachusetts General Hospital, Boston, MA 02114 USA

## Abstract

Mechanical transformation of tissue is not merely a symptom but a decisive driver in pathological processes. Comprising intricate network of cells, fibrillar proteins, and interstitial fluid, tissues exhibit distinct solid- (elastic) and liquid-like (viscous) behaviours that span a wide band of frequencies. Yet, characterization of wideband viscoelastic behaviour in whole tissue has not been investigated, leaving a vast knowledge gap in the higher frequency range that is linked to fundamental intracellular processes and microstructural dynamics. Here, we present wideband Speckle rHEologicAl spectRoScopy (SHEARS) to address this need. We demonstrate, for the first time, analysis of frequency-dependent elastic and viscous moduli up to the sub-MHz regime in biomimetic scaffolds and tissue specimens of blood clots, breast tumours, and bone. By capturing previously inaccessible viscoelastic behaviour across the wide frequency spectrum, our approach provides distinct and comprehensive mechanical signatures of tissues that may provide new mechanobiological insights and inform novel disease prognostication.

## Introduction

Mechanical properties of tissue play an important role in regulating cellular functions and driving disease processes^1-3^. Aberrant mechanical remodelling is implicated in a broad spectrum of pathologies including the onset and progression of neoplasms^4-6^, haematological disorders^7,8^, cardiovascular diseases^9-11^, fibro-proliferative disorders^12^ and several orthopaedic conditions^13^. For instance, alterations in tissue stiffness, a frequent consequence of desmoplastic reaction has been linked to malignancy and chemo-resistance in solid tumours^14-18^. Compromised bone strength is associated with diminished bone density and may portend a higher risk of osteoporotic fractures^19,20^. Coagulopathy is associated with modulation in the mechanical properties and stability of blood clots^21^, motivating the development of mechanics-based point-of-care diagnostic devices for haemostasis management^22^. Meanwhile, optical, ultrasound and magnetic resonance elastography techniques are utilized for the clinical management of liver fibrosis, breast lesions, multiple sclerosis, and other tissue pathologies^23-25^.

The evidence that mechanical factors are decisive participants in disease pathogenesis is unequivocal. Current insights, however, have largely relied on a single mechanical descriptor, elasticity, that is inadequate for capturing the full complexity of mechanical cues that regulate mechanobiological processes. Tissue is composed of a complex network of cells and fibrillar proteins, surrounded by interstitial fluid that contains 90% water. The structural hierarchy within the tissue that spans nano-to millimetre length scales is accompanied by a comparably vast breadth of relaxation time scales^26-28^. These intricate structure and composition of tissue yield both elastic (*G*′) and viscous (*G*″) behaviours that together contribute to the complex shear (viscoelastic) modulus, *G*\*(*ω*)=*G*′(*ω*)+*iG*″(*ω*), modulated over a wide range of angular frequency, *ω*. Thus, living cells sense both elastic and viscous mechanical cues at distinct *ω*-frequencies depending on how fast they interact with their microenvironment^29-31^. Notably, cells respond to viscous dissipation in the ECM via intracellular signalling in ways not explained by changes in elasticity alone^32-35^. In 3D cell culture models, cancer cell invasion is simultaneously accompanied by two seemingly opposing mechanical changes in the ECM: stiffening and ‘liquidisation’, each dominating at different frequency regimes^36^. Therefore, approaches to measure frequency-dependent elastic and viscous moduli over a broad spectrum—that is, wideband micromechanical spectroscopy approaches—are needed to comprehensively capture distinct viscoelastic behaviours in tissues and other complex biological materials.

Seminal microrheological studies in homogeneous materials have revealed distinct viscoelastic signatures that span a frequency range of over 6-decades or beyond^37,38^. Some hydrogels such as polyacrylamide may appear linear elastic under classical macrorheology (e.g., with a rheometer tool), but exhibit distinct frequency-dependent viscoelastic behaviour over multiple decades of frequency under microrheology^39-41^. Rich frequency-dependent viscoelastic behaviours-most notable being the high-frequency power scaling laws in the kHz-MHz regime-have since been observed in cells^42-45^, biofluids^46^, and ECM constructs^47-49^. Recent microrheological studies in reconstituted ECM and *in vitro* cell culture point to novel mechanobiological insights that are only accessible in the kHz-MHz regime. However, wideband micromechanical spectroscopy has never been investigated in whole (intact) tissue. Study in this regime has the potential to provide new sources of micromechanical contrast for improved disease prognostication and generate new insights in the field of mechanobiology that has thus far relied on a single elastic modulus at a low frequency.

Existing techniques do not support frequency-dependent measurement of elasticity and viscosity up to the MHz regime in whole tissue^50^. Conventional rheometry can only provide bulk measurement at very low frequencies, typically over a few Hz^51^. Although conventional microrheology techniques, based on particle tracking^52,53^, dynamic light scattering (DLS)^37,46^, diffusing wave spectroscopy (DWS)^38,54^, and optical manipulation^45,55-57^, are capable of measurement over a wide frequency range, they are not applicable in whole tissue due to the reliance on exogenous probe particles. Furthermore, particle tracking approaches are typically limited to highly compliant samples (shear modulus <100 Pa) while DLS-based techniques are limited to transparent diluted samples (i.e., single-scattering limit). Among techniques that can be applied in whole tissue, optical coherence elastography (OCE) typically provides quasi-static measurement of elasticity or dynamic measurement over limited frequencies (<10 kHz)^58-60^. Meanwhile, Brillouin microscopy provides longitudinal modulus in the GHz range at high spatial resolution, but the interpretation of this modulus in relation to widely recognized Young’s or shear modulus remains a challenge^61^. Thus, there is a large knowledge gap in the kHz-MHz range, a regime in which the mechanical behaviour of tissue remains largely unknown; yet, one that is linked to fundamental intracellular processes and microstructural dynamics.

Here, we present Speckle rHEologicAl spectRoScopy (SHEARS) to address the need for frequency-dependent viscoelastic analysis in whole tissue up to the sub-MHz regime. We have previously shown that speckle formed by the interference of backscattered laser illumination is sensitive to the natural thermal motion of native light scattering structures in biological materials, and thus, can be leveraged to provide passive microrheological analysis in an entirely all-optical, non-invasive, and noncontact manner^62-65^. Based on this principle, we developed an approach to map mechanical properties of whole tissue with high spatial resolution and demonstrated measurement of shear modulus magnitude, |*G*\*|, over a frequency range of *ω*=1–250 rad/s in various types of biological specimens^65-71^. However, these prior studies did not realize the wideband frequency-dependent elastic, *G*′(*ω*), and viscous, *G*″(*ω*), spectra that distinctly contributed to the complete viscoelastic behaviour of tissue. With SHEARS, we demonstrate, for the first time, wideband micromechanical spectroscopy of shear elastic and viscous moduli up to the sub-MHz frequency regime in biomimetic scaffolds and whole tissues. We analyse the wideband viscoelastic spectra of fibrin constructs, whole blood clots, breast tumours, and bone. We show that distinct frequency-dependent elastic and viscous signatures exist over multiple frequency regimes up to the sub-MHz range. Our results demonstrate an unprecedented dynamic range in both the mechanical properties of the specimens (fluid to bone) and the measurement frequencies (1–10^5^ rad/s) simultaneously in a single platform.

## Results

### Principle of wideband SHEARS

SHEARS is an advancement over laser speckle rheology approaches previously developed by our group^62-65^. Biological materials such as tissue are composed of numerous microscopic light scattering structures that are susceptible to naturally occurring thermal (Brownian) displacements, the magnitude and frequencies of which are governed by the local viscoelastic behaviour of the microenvironment. Consequently, scattering structures in softer cellular or adipose tissues may exhibit larger and more rapid displacements compared to stiffer fibrous regions. Upon illumination by a coherent beam, light scattered by these dynamic structures interfere to form a temporally fluctuating speckle pattern. By recording a time series of speckle patterns and analysing its magnitude and rate of fluctuation, the frequency-dependent viscoelastic behaviour of the material can be reconstructed. Unlike conventional DLS and DWS approaches, laser speckle rheology is not restricted to either transparent diluted sample (for DLS) or highly concentrated turbid suspension (for DWS) that require incorporating specific amount of exogenous scattering particles. Rather, it directly exploits displacements of native scatterers that are already present in the sample and navigates a range of intrinsic tissue scattering properties that lies between the two scattering limits. Based on this principle, measurement of |*G*\*| at low frequency (*ω*<250 rad/s) has been validated and applied in hydrogels^65^, blood^67-69^, atherosclerotic plaques^66,70^, and breast tumours^71^. The new wideband SHEARS approach detailed here extends the capabilities of laser speckle rheology to permit measurement of the full complex modulus as well as the distinct contributions of elastic and viscous moduli, *G*′(*ω*) and *G*″(*ω*), over *ω*∼1–10^5^ rad/s in whole tissue.

The highest frequency accessible by SHEARS is physically limited by the speckle acquisition frame rate, which directly determines the smallest time scale at which the rate of speckle fluctuation can be measured. We utilize a high-speed CMOS camera (Photron, Mini AX200 type 900k) that can support a frame rate up to 540 kHz with a sensor ROI of 128×32 pixels. The optical system is designed to achieve a pixel-to-speckle ratio of 3.5 pixels/speckles, ensuring sufficient spatial sampling to capture fully developed speckle, while also maximizing the number of individual speckles captured within the small sensor ROI. Speckle time series is acquired in a 180° backscattered configuration at perpendicular polarization to the illumination (Fig. 1a, see Methods for detailed description of the optical system and speckle acquisition procedure). Speckle fluctuation is evaluated by computing the ensemble-averaged intensity autocorrelation function, *g*_2_(*t*), from which the time-dependent mean square displacement (MSD) of the scatterers is obtained according to equations (1) and (2) in Methods^62,63,65^. Notably, equation (2) is an empirical approximation of the DWS formulation that allows SHEAR to account for an arbitrary set of optical properties in each sample^62,63^. An example of *g*_2_(*t*) curve and time-dependent MSD measured in a fibrin hydrogel are shown in Fig. 1b (see Methods for fibrin sample preparation).

**Fig. 1.**
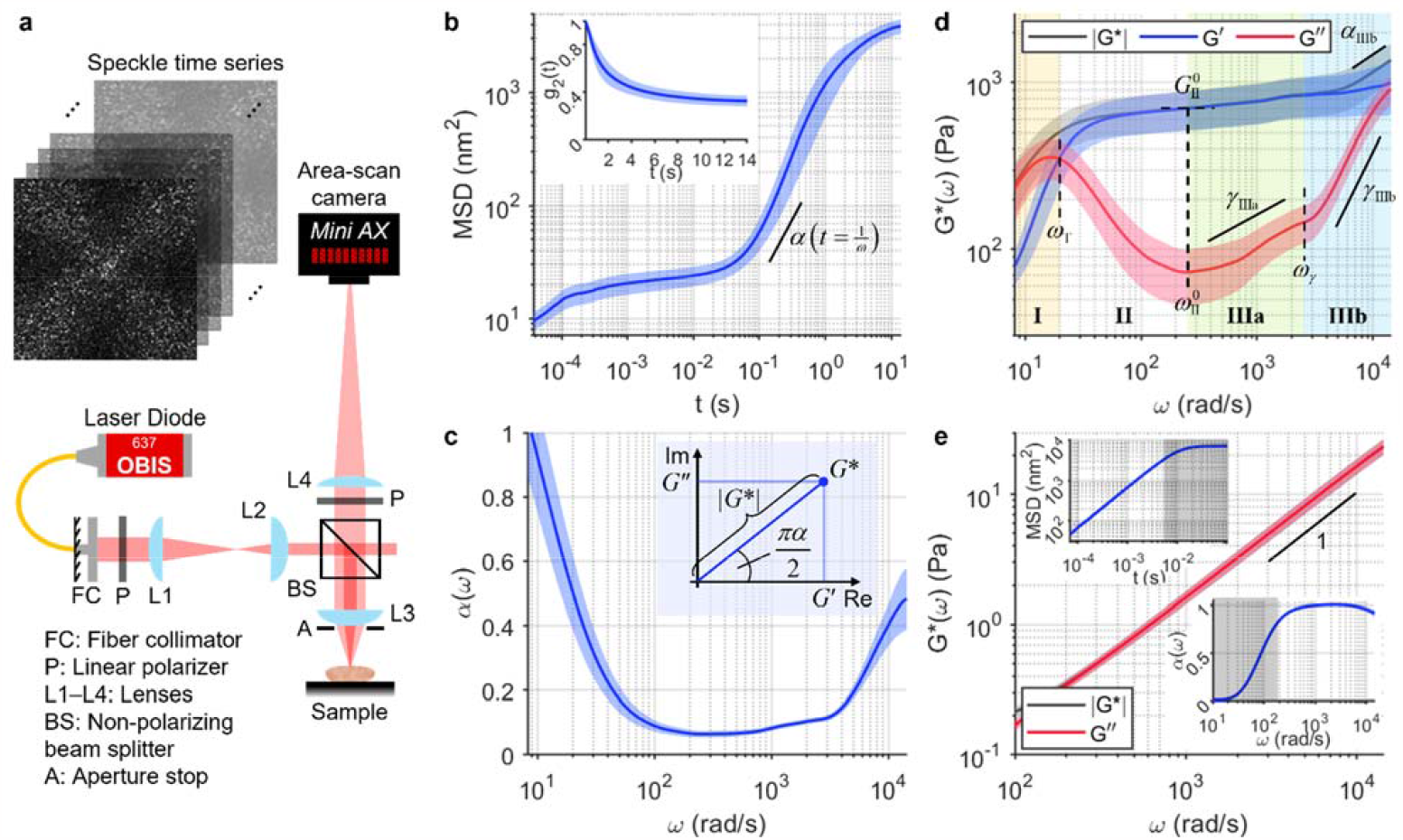
Wideband micromechanical spectroscopy with SHEARS. **a** Optical setup of the SHEARS system. **b** Time-dependent MSD of polymerised fibrin construct computed from the intensity autocorrelation (inset) of the acquired speckle time series. **c** Frequency-dependent *α*(*ω*) obtained from the log-log derivative of the MSD in **b**. Inset illustrates the role of *α* in determining the relative contribution of *G*′ and *G*″ in the complex modulus, *G*\*=*G*′+*iG*″. **d** Frequency-dependent |*G*\*(*ω*)| (black), *G*′(*ω*) (blue), and *G*″(*ω*) (red) of fibrin construct computed from MSD and *α* in **b** and **c**. Values of all annotated spectroscopic parameters are tabulated in Supplementary Table 1. **e** |*G*\*(*ω*)| (black) and *G*″(*ω*) (red) of pure fibrinogen solution containing 3-μm diameter polystyrene particles, showing linearly increasing *G*\*(*ω*)=*G*″(*ω*) indicated by a log-log slope of 1. Insets show the measured MSD and *α*; shaded areas indicate time scales over which the speckle pattern has completely decorrelated due to the fast dynamics in aqueous solution. In **b**– **e**, solid curve and shaded outline represent mean±standard deviation of *N*=9 measurements, respectively.

In order to reconstruct both the elastic, *G*′(*ω*), and viscous, *G*″(*ω*), moduli, the log-log derivative of the MSD w.r.t time (slope annotated on Fig. 1b) is computed to obtain the frequency-dependent power scaling law, *α*(*ω*), where MSD∼*t*^*α*(*ω*)^ at *t*=1/*ω* (Fig. 1c). While the MSD itself is inversely proportional to the magnitude |*G*\*|, its power scaling law *α* determines the phase angle ∠*G*\* that governs the relative contribution of the real, *G*′, and imaginary, *G*″, parts of the complex modulus (inset of Fig. 1c); equation (3) in Methods provides the full expression^72-74^. Here, *α*=0 indicates a purely elastic solid-like behaviour (i.e., |*G*\*|=*G*′) whereas *α*=1 indicates a purely viscous fluid-like behaviour (i.e., |*G*\*|=*G*″). Meanwhile, viscoelastic materials exhibit frequency-dependent *α*(*ω*) that varies between the two limits (i.e., 0<*α*<1), where *α*=0.5 corresponds to *G*′=*G*″ and deviation in either direction indicates more elasticity-dominant (*α*<0.5, *G*′>*G*″) or viscosity-dominant (*α*>0.5, *G*′<*G*″) behaviour. For the fibrin construct, *α*(*ω*) decreases from *α*∼1 at *ω*≤10 rad/s to *α*∼0.1 at 100≤*ω*<3,000 rad/s then approaches *α*∼0.5 at *ω*>10^4^ rad/s (Fig. 1c), giving rise to frequency-dependent *G*′(*ω*) and *G*″(*ω*) spectra (Fig. 1d). In contrast, a solution of unpolymerized fibrinogen molecules (fibrin precursors) exhibits a linearly increasing MSD and a constant *α*(*ω*)=1 (insets of Fig. 1e), resulting in a purely viscous behaviour with |*G*\*|=*G*″, where *G*″ increases linearly with *ω* as expected of a viscous fluid (Fig. 1e).

### Frequency-dependent viscoelastic behaviour of fibrous polymer construct

We measured *G*′(*ω*) and *G*″(*ω*) of polymerised fibrin constructs to examine the frequency-dependent viscoelastic behaviour of a typical fibrous ECM (Fig. 1d). Fibrin was chosen as an example for its clinical relevance in haematology and wound healing. Wideband SHEARS up to *ω*>10^4^ rad/s reveals 4 viscoelastic regimes across the frequency spectrum. In Regime I, the viscoelastic behaviour of the fibrin construct is dominated by viscous contribution with *G*″>*G*′. The low-frequency fluid-like behaviour is attributed to the relaxation dynamics of the hydrogel construct as a whole^75,76^. As the frequency increases, *G*′ gradually approaches *G*″ until they crossover (i.e., *α*=0.5) at the fluid-to-solid transition frequency, *ω*_T_=20±3 rad/s. In Regime II, the fibrin construct reaches its most elastic behaviour with *α*<0.1 (Fig. 1c) as *G*″ decreases to a local minimum at frequency *ω*^0^_II_=300±100 rad/s while *G*′ reaches the ‘elastic plateau modulus’ of *G*^0^_II_=700±200 Pa. The elastic plateau is governed by the elasticity of the fibrin fiber network. Here, the elastic plateau extends for ∼2 decades with both *G*′ and |*G*\*| remaining relatively constant. In Regime III, hidden beneath the elastic plateau, *G*″ increases following a power scaling law *G*″∼*ω*^*γ*^. Two distinct scaling laws can be observed: first with the exponent of *γ*_IIIa_=0.4±0.1, then transitions to *γ*_IIIb_=1.2±0.1 at frequency *ω*_γ_=17×10^3^±2×10^3^ rad/s. Within Regime IIIb, the increasing contribution of *G*″(*ω*) causes the overall modulus magnitude to also follow a power scaling law |*G*\*|∼*ω*^*α*^ with an exponent of *α*_IIIb_=0.50±0.09. The shift from |*G*\*| plateau to high-frequency power scaling can be interpreted as the transition from the network-level elastic behaviour at intermediate frequencies to the single filament-level fluctuation at higher frequencies^46,75,76^. The power scaling exponent of *α*∼0.5 is consistent with the bending fluctuation of a Rouse flexible polymer^75,76^.

Values of all spectroscopic parameters annotated in Fig. 1 are summarised in Supplementary Table 1. These wideband spectroscopic parameters together describe the intricate frequency-dependent elastic and viscous behaviour of polymerised fibrin, a fibrillar protein commonly found in biological tissue. In the following sections, we show that the spectroscopic analysis described above can be applied to a wide range of clinical specimens to reveal distinct mechanical signatures in tissues such as whole blood clots, breast tumours, and bone.

### Wideband micromechanical spectroscopy of blood clot viscoelasticity

Blood clots are essentially polymerised fibrin scaffolds that contain various other cellular components such as platelets and red blood cells (RBC). Mechanical properties of blood clots have emerged as an important player in the management of thrombotic and bleeding complications in the clinical settings^68,69,77^. We obtained two clinical blood samples from the MGH Core Laboratory (MGH IRB#2017P000419), each presenting high (hereafter ‘high-FIB’) and low (hereafter ‘low-FIB’) fibrinogen content associated with potential thrombotic and bleeding risks, respectively. Wideband SHEARS was performed in whole blood before (Fig. 2, inset) and after (Fig. 2) clot initiation by Kaolin and CaCl_2_. Values of all spectroscopic parameters annotated in Fig. 2 are summarised in Supplementary Table 1.

**Fig. 2.**
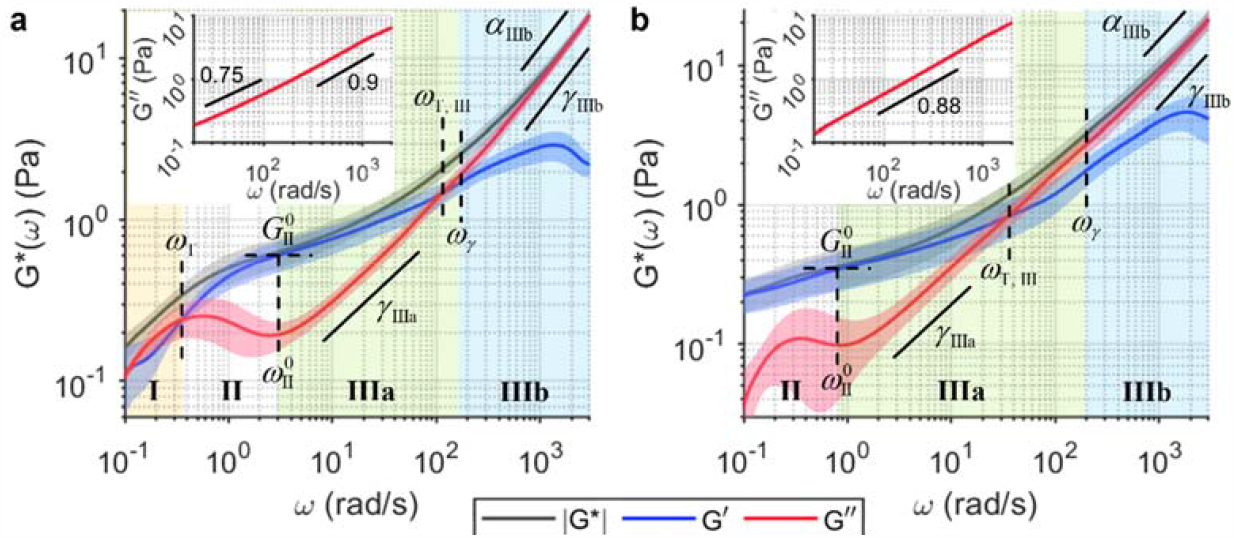
Wideband viscoelastic spectra of clinical blood clots. **a, b** Frequency-dependent |*G*\*(*ω*)| (black), *G*′(*ω*) (blue), and *G*″(*ω*) (red) of clots formed by whole blood samples with fibrinogen contents at the upper (‘high-FIB’, 5.15 mg/mL) and lower (‘low-FIB’, 2.12 mg/mL) limits of the normal range, respectively. Solid curve and shaded outline represent mean±standard deviation of *N*=5 measurements. Values of all annotated spectroscopic parameters are tabulated in Supplementary Table 1. Insets show *G*″(*ω*) in pure whole blood prior to clot initiation. Annotated numbers indicate the power scaling law of *G*″(*ω*).

The viscoelastic behaviour of high-FIB clot is similar to that of the purified fibrin construct in Regimes I and II, but with both the fluid-to-solid transition and the elastic plateau occurring at lower frequencies of *ω*_T_=0.36±0.09 rad/s and *ω*^0^_II_=3.1±0.7 rad/s, respectively (Fig. 2a). Furthermore, the elastic plateau modulus is lower at *G*^0^_II_=0.6±0.2 Pa and extends for less than a decade, indicating lower network elasticity and an overall less solid-like behaviour compared to the purified fibrin in Fig. 1d. In Regime III, *G*″(*ω*) power scaling has the exponent of *γ*_IIIa_=0.6±0.2 that transitions to *γ*_IIIb_=0.83±0.01 at *ω*_γ_=160±30 rad/s, suggesting that the viscous behaviour approaches that of a purely viscous fluid with *γ*→1 (i.e., *G*″ increases linearly with *ω*) at higher frequencies. A unique behaviour in whole blood clots compared to purified fibrin is the second crossover between *G*′ and *G*″ at *ω*_T,III_=110±40 rad/s—this time, a transition back to a more viscosity-dominant behaviour with *G*″ exceeding *G*′. With *G*″ dominating the viscoelastic behaviour after this transition, |*G*\*(*ω*)| follows the same power scaling law with exponent *α*_IIIb_=*γ*_IIIb_. In comparison, the low-FIB clot has a lower elastic plateau modulus of *G*^0^_II_ =0.3±0.2 Pa that occurs at a frequency of *ω*^0^_II_ =0.8±0.7 rad/s (Fig. 2b). Furthermore, the solid-to-fluid transition in Regime III also occurs at a much lower frequency of *ω*_T,III_=40±30 rad/s compared to high-FIB. Thus, the low-FIB clot not only has lower elasticity (i.e., more compliant) than the high-FIB clot, but also exhibits more *G*″-dominant behaviour across a wider range of frequencies, likely due to the lower availability of fibrinogen protein to form a stable fibrin network.

Evidently, the viscoelastic spectra of clots formed by whole blood (Fig. 2) are markedly different from those of the purified fibrin construct (Fig. 1d), despite both being primarily composed of fibrin network. The discrepancies are likely due to the presence of other microscopic components in whole blood such as platelets and RBCs, resulting in altered structure of the fibrin fibres compared to the purified scaffold. The complexity of whole blood can also be appreciated from the viscosity measurement prior to clot initiation. Unlike in solution of unpolymerized fibrinogen molecules (Fig. 1e), *G*″ does not increase linearly with frequency (i.e., *α*=1) but instead follows a power law with *α*(*ω*) in the range of 0.75–0.9 (insets of Fig. 2). Furthermore, both whole blood samples are more viscous, with average viscosity of 4.5±0.2 mPa·s (high-FIB) and 4.6±0.1 mPa·s (low-FIB) at *ω*>200 rad/s, compared to the pure fibrinogen solution with viscosity of 1.43±0.06 mPa·s.

### Profiling wideband viscoelastic signatures in breast cancer

Breast tumours are mechanically heterogeneous, both across different molecular subtypes and within a single specimen. We have previously shown that the cellular tumour epithelium tends to have lower |*G*\*| than the fibrous stroma, with tumours of more aggressive molecular subtypes generally exhibiting sharper |*G*\*| gradient across the tumour-stroma interface^71^. We performed wideband SHEARS on three freshly excised breast specimens from different patients (MGH IRB#2011P000301): benign breast tissue from a patient with breast cancer (Fig. 3a, b), residual tumour after treatment with neoadjuvant chemotherapy (Fig. 3c–e), and untreated invasive breast carcinoma (Fig. 3f–j). Across the three specimens, we observe four different types of frequency-dependent viscoelastic behaviour corresponding to benign fibrous breast tissue (Fig. 3b), adipose (Fig. 3e), fibrous tumour stroma (Fig. 3d, g, i), and cellular tumour epithelium (Fig. 3h, i). Values of all spectroscopic parameters annotated in Fig. 3 are summarised in Supplementary Table 1.

**Fig. 3.**
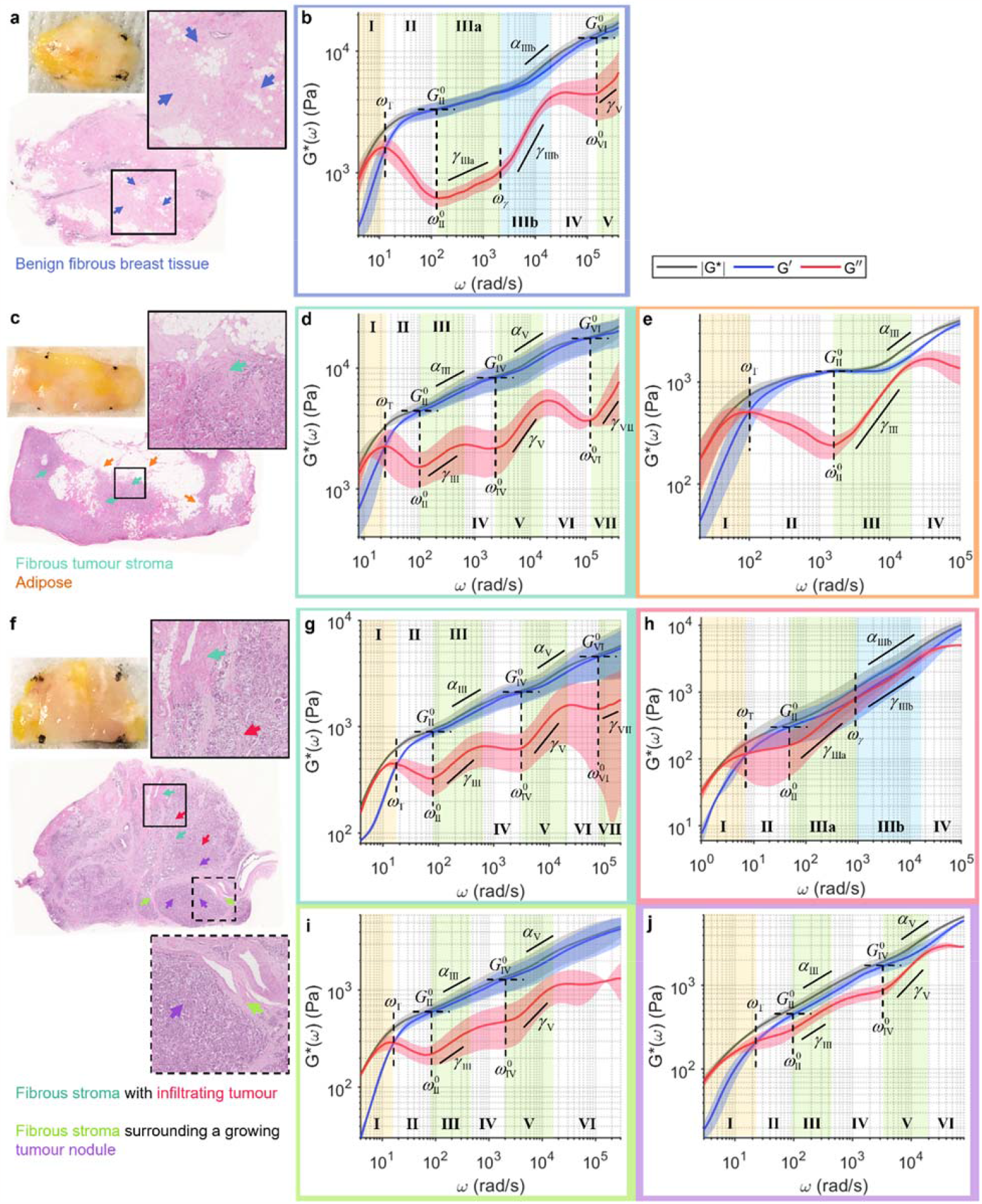
Wideband viscoelastic spectra of the breast tumour microenvironments. Gross photo, H&E slide, and frequency-dependent |*G*\*(*ω*)| (black), *G*′(*ω*) (blue), and *G*″(*ω*) (red) of **a, b** benign breast tissue, **c**–**e** treated invasive carcinoma, and **f**–**j** untreated invasive carcinoma. Solid curve and shaded outline represent mean±standard deviation of measurements at locations indicated by coloured arrows on H&E. Arrows and plot boxes are colour-coded for benign fibrous breast tissue (blue), adipose (orange), tumour stroma (dark green, light green), and cellular tumour epithelium (purple, red). Values of all annotated spectroscopic parameters are tabulated in Supplementary Table 1.

In benign fibrous breast tissue (Fig. 3a), the wideband viscoelastic behaviour is rather reminiscent of that of the purified fibrin constructs in Fig. 1d, except with the moduli magnitude being overall higher (Fig. 3b). Notably, two distinct power scaling laws are also observed for *G*″(*ω*) in Regime III, with both exponents being lower than those of fibrin (*γ*_IIIa_=0.20±0.03, *γ*_IIIb_=0.73±0.05) but transitioning at similar frequency of *ω*_γ_=2000±200 rad/s. Similarly, the increasing contribution of *G*″(*ω*) in Regime IIIb also results in the high-frequency power law behaviour of |*G*\*(*ω*)| with the exponent of *α*_IIIb_=0.33±0.05. Taking the measurement up to *ω*>10^5^ rad/s reveals two additional frequency regimes. Regimes IV-V exhibit similar behaviours to Regimes II-III, with *G*″ reaching another local minimum at frequency *ω*^0^_IV_=2×10^5^±1×10^5^ rad/s accompanied by its own elastic plateau modulus of *G*^0^_IV_=13±2 kPa, followed by a power scaling law with exponent of *γ*_V_=0.5±0.1.

Invasive ductal carcinoma after neoadjuvant (presurgical) systemic treatment displays varying levels of cellularity, with interspersed tumour cell clusters within the fibrous tumour stroma and areas of normal adipose (Fig. 3c). Unlike the benign fibrous breast tissue, the fibrous tumour stroma (green arrows in Fig. 3c) exhibits a characteristic undulation pattern in its viscous behaviour (Fig. 3d), where the viscoelastic spectra repeatedly reach a new elastic plateau (Regimes II, IV, and VI) followed by a *G*″(*ω*) power scaling (Regimes III, V, and VII), each obeying only a single power law (i.e., *γ*_IIIb_ in Fig. 3b is not observed). Notably, higher elastic plateau modulus is observed at increasing frequency regimes (*G*^0^_II_=4.6±0.9, *G*^0^_IV_=8±2, *G*^0^_VI_=18±5 kPa), indicating decreased compliance as a function of frequency. Interestingly, the power scaling laws also follow a similar increasing trend for both *G*″ (*γ*_III_=0.33±0.09, *γ*_V_=0.55±0.06, *γ*_VII_=0.6±0.3) and |*G*\*| (*α*_III_=0.22±0.06, *α*_V_=0.26±0.01). An increase in the high-frequency power law exponent may be interpreted as a decrease in the filament flexibility of the fibrillar structures^75,76^. Meanwhile, the viscoelastic spectra of adipose tissue (orange arrows in Fig. 3c) exhibit only one elastic plateau and one *G*″(*ω*) power scaling regime (Fig. 3e). The elastic plateau occurs at a higher frequency of *ω*^0^_II_=1,500±200 rad/s with a lower plateau modulus of *G*^0^_II_=1.27±0.06 kPa compared to the fibrous stroma. However, the high-frequency power scaling of both *G*″(*ω*) and |*G*\*(*ω*)| are steeper with the exponents of *γ*_III_=0.8±0.1 and *α*_III_=0.44±0.05, respectively.

Compared to the treated sample, the untreated invasive carcinoma displays well-delineated regions of fibrous stroma and cellular tumour epithelium (Fig. 3f). The fibrous stroma with tumour infiltration (dark green arrows in Fig. 3f, upper inset) exhibits viscoelastic behaviour that is remarkably similar to that of the stroma in the treated tumour in Fig. 3d, with the characteristic undulation pattern in *G*″(*ω*) (Fig. 3g). Notably, although the elastic plateau moduli are lower (*G*^0^_II_ =0.9±0.09, *G*^0^_IV_ =2.1±0.4, *G*^0^_VI_ =5±2 kPa), the power scaling exponents for both *G*″(*ω*) (*γ*_III_ =0.44±0.03, *γ*_V_=0.58±0.06, *γ*_VII_=0.35±0.3) and |*G*\*(*ω*)| (*α*_III_=0.27±0.05, *α*_V_=0.28±0.07) are in the same range as those of the treated tumour. Likewise, the fibrous tumour stroma surrounding a growing nodule of tumour cells (light green arrows in Fig. 3f, lower inset) also exhibits characteristic undulation in the viscous behaviour and power scaling exponents in the same range of *α*∼0.2–0.3 (Fig. 3i). In comparison, the cell-dense nodule of tumour (purple arrows in Fig. 3f) exhibits noticeably more viscous behaviour, where *G*″ lies close to *G*′ throughout the spectra (Fig. 3j). The higher contribution of *G*″ to the overall modulus is also reflected in the power scaling exponents of |*G*\*(*ω*)|, with *α* approaching 0.5 (*α*_III_=0.40±0.01, *α*_V_=0.45±0.07). The viscous behaviour is even more dominant in the tumour that is not contained in a nodule but infiltrates the fibrous stroma (red arrows in Fig. 3f), where *G*″ completely overlaps *G*′, resulting in a high frequency |*G*\*(*ω*)| power scaling law with *α*_IIIb_=0.51±0.09 that extends over a decade (Fig. 3h). The relatively more fluid-like behaviour of the infiltrating tumour supports the model of cancer invasion based on cell jamming theory, where the invading tumour appears liquidised as it ‘unjams’ from its solid tumour state^78-80^.

Our results show that the breast tumour microenvironment is an extremely heterogeneous micromechanical landscape, not only in quasi-static elasticity^81-84^ or low-frequency viscoelastic modulus^71^, but even more remarkably so in the frequency-dependent elastic and viscous behaviours over a wide frequency spectrum. Notably, although all tissues in Fig. 3 behave similarly at *ω*<100 rad/s (the limited frequency range in which many mechanobiological studies have been based), they exhibit vastly distinct viscous behaviour at the higher frequency regimes. These wideband viscoelastic spectra may lead to further insights on the multifaceted mechanical transformation in breast cancer and provide new sources of contrast for disease prognostication.

### Revealing wideband frequency-dependent viscoelastic behaviours of bone

Although bone mineral density (BMD) has long served as a diagnostic marker for bone diseases, it is now accepted that BMD is an incomplete index of bone fragility and does not adequately predict fracture risk across disease states^19^. Furthermore, there is a growing appreciation for the role of structural water on the mechanical integrity of bone in aging and diseases^85,86^. However, mechanical properties of bone are still typically given by elastic moduli, which exhibits little frequency dependence at the macroscale^13^. Microrheology of bone remains largely unexplored in orthopaedic conditions. Leveraging the large dynamic range of wideband SHEARS, not only in the frequency but also in the measurable viscoelastic modulus, we reveal for the first time, frequency-dependent elastic and viscous behaviours of bone across a wide range of frequency (Fig. 4). Bovine rib bone was cross-sectioned (Fig. 4a) to obtain wideband viscoelastic spectra in the dense cortical bone at the outer shell (Fig. 4b) and the ‘spongy’ trabecular bone at the centre (Fig. 4c). Values of all spectroscopic parameters annotated in Fig. 4 are summarised in Supplementary Table 1.

**Fig. 4.**
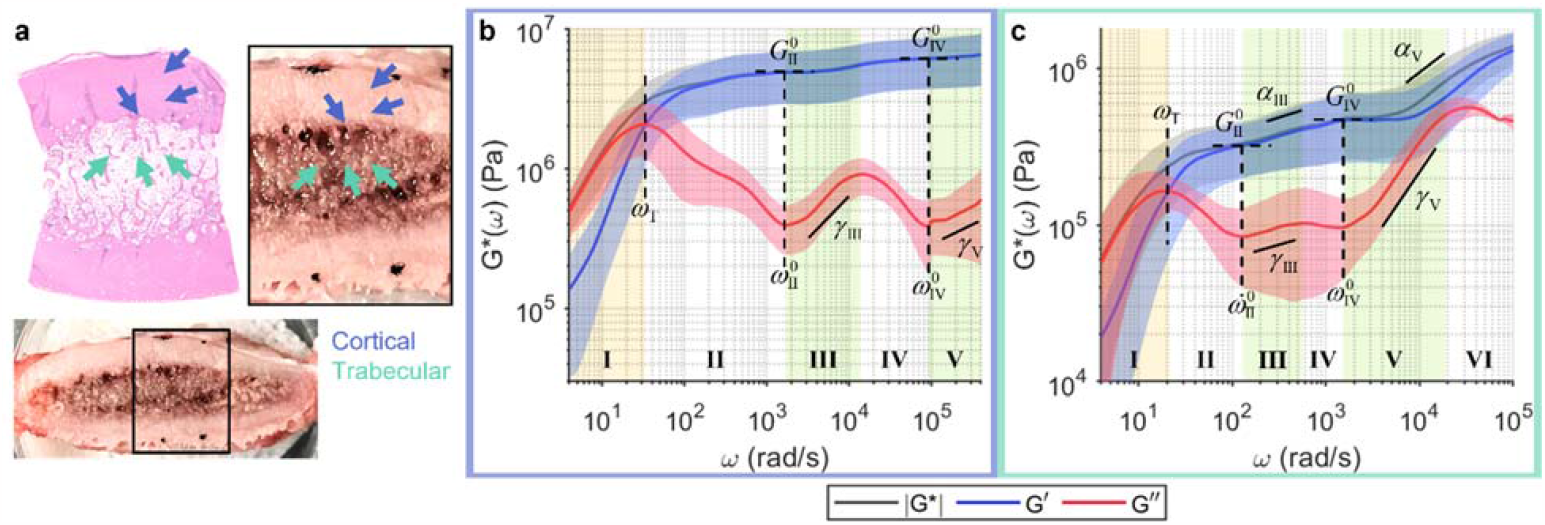
Wideband viscoelastic spectra of compact and trabecular bones. **a** Gross photo and H&E slide of bovine rib bone cross-section. **b, c** Frequency-dependent |*G*\*(*ω*)| (black), *G*′(*ω*) (blue), and *G*″(*ω*) (red) of cortical (blue) and trabecular (green) bones indicated by coloured arrows in **a**, respectively. Solid curve and shaded outline represent mean±standard deviation of measurements at locations indicated by coloured arrows on H&E. Values of all annotated spectroscopic parameters are tabulated in Supplementary Table 1.

At the microscale, both cortical and trabecular bones exhibit frequency-dependent elastic and viscous behaviours not unlike those of soft tissues. The cortical bone, which is composed of densely packed osteons with aligned fibrous collagen matrix, displays an undulation pattern in the viscous behaviour that is characteristic of the fibrous stroma of breast tumours (Fig. 4b). Compared to the fibrous tumour stroma, the undulation of *G*″ begins (i.e., the first local minimum) at a much higher frequency of *ω*^0^_II_=1,720±70 rad/s compared to *ω*^0^_II_∼100 rad/s in Fig. 3d, g, i. In addition, although *G*″(*ω*) follows a similar power scaling behaviour (*γ*_III_=0.56±0.09, *γ*_V_=0.2±0.2), |*G*\*(*ω*)| remains roughly constant over the measured frequency range (i.e., *α*→0) with elevated elastic plateau moduli of *G*^0^_II_=5±2 and *G*^0^_IV_=6±2 MPa. Evidently, the wideband viscoelastic behaviour of the cortical bone is more predominantly elastic than that of the fibrous tumour stroma even though both are highly fibrous tissues with interspersed cells (tumours and osteocytes), likely owing to the calcification and mineral contents in the bone. Meanwhile, the trabecular bone, which is composed of less densely arranged trabeculae surrounding soft tissue components such as marrow and blood vessels, exhibit lower elastic plateau moduli (*G*^0^_II_=0.3±0.1, *G*^0^_IV_=0.5±0.2 MPa) compared to the cortical bone (Fig. 4c). Furthermore, the frequency-dependent elastic and viscous behaviours also bare more resemblance to those of other fibrous soft tissues and scaffolds, with both fluid-to-solid transition and elastic plateau occurring at the same frequency ranges of *ω*_T_∼20 rad/s and *ω*^0^_II_∼100 rad/s, respectively. Moreover, the power scaling behaviours of both *G*″(*ω*) (*γ*_III_=0.1±0.1, *γ*_V_=0.78±0.06) and |*G*\*(*ω*)| (*α*_V_=0.44±0.07) are particularly similar to those of the benign fibrous breast tissue in Fig. 3b.

Although microrheology has traditionally been reserved for the study of complex fluids and soft materials, our results show that bone also exhibits rich frequency-dependent viscoelastic behaviour when probed at the microscale. With increasing attention to not only solid materials (e.g., collagen and minerals) but also structural water in bone^85,86^, examination of both elastic and viscous behaviours of the bone across a wide range of frequency scales is likely to offer new insights on the microstructural mechanisms underlying orthopaedic conditions.

## Discussion

In this work, we harness the interactions of coherent laser light and thermally-driven micromechanical fluctuations of native scattering structures in biomaterial constituents to obtain ‘viscoelastic fingerprints’ over a wide frequency spectrum in tissues, biofluids, and biomimetic scaffolds. These wideband viscoelastic spectra can enable characterization of multiscale mechanics, support biophysical studies of microstructural dynamics in cells and ECM, and provide new means to investigate the mechanobiology of diseases.

From the rheological perspective, wideband micromechanical spectroscopy over a broad range of frequency scales inherently interrogates a comparably wide range of length scales within the structural hierarchy of the material. Cells and extracellular networks that form the tissue at the mesoscale are composed of building blocks at length scales of micrometres down to nanometres and beyond, where structures of smaller characteristic length scales are generally associated with faster dynamical behaviours^26-28^. Thus, viscoelastic moduli that span the lower to higher frequency regimes correspond to the mechanical behaviour of the whole tissue structure (collective network dynamics) down to the individual constituents (single-filament dynamics) that compose the tissue^46,76,87^, despite the fixed spatial resolution of the optical system. Indeed, at lower frequencies (*ω*<30 rad/s), our prior studies have demonstrated agreement between *G*\*(*ω*) measured with laser speckle microrheology approaches and conventional macrorheology in aqueous solution^62,63^, homogeneous hydrogels^65^, and breast tissues^71^. Here, the full wideband viscoelastic spectra in fibrin constructs (Fig. 1d) show that *G*\*(*ω*) most closely corresponds to macrorheology measurement at the elastic plateau (*ω*=*ω*^0^_II_), where the viscous modulus is approximately 10% of the elastic modulus^29^. This is consistent with the interpretation that the elastic plateau is governed by collective network elasticity^46,75,76^. Beyond the plateau, wideband SHEARS provides additional rheological information that is inaccessible to conventional macrorheology.

From the biophysical perspective, wideband micromechanical spectroscopy reveals characteristic frequency-dependent behaviours that facilitate physics-based interpretation of biomaterial properties. Despite vastly diverse spectral signatures across different types of tissues, we show that the wideband spectra can be divided into smaller frequency regimes, within which the frequency-dependent behaviour follows characteristic patterns that have been previously predicted by various biophysical models. Taking the fibrin construct (Fig. 1d) as an example, the viscoelastic behaviour is viscosity-dominant (*G*″>*G*′) at the lowest frequencies before the crossover of *G*′ and *G*″ indicates the shift to elasticity-dominant regime (Regime I). Then, *G*″ decreases to a minimum at *ω*=*ω*^0^ while *G*′ remains relatively constant, resulting in |*G*\*|∼*G*^0^, the elastic plateau modulus (Regime II). These behaviours are well described by the Maxwell model^88,89^. As *G*′ remains plateaued, *G*″ rises from its minimum following power scaling laws *G*″∼*ω*^*γ*^ (Regime IIIa and IIIb). Eventually, the increasing viscous contribution causes the modulus magnitude to deviate from the plateau and follow a power scaling law |*G*\*|∼ *ω*^*α*^ at higher frequencies (Regime IIIb). These behaviours are predicted by various models of Rouse polymer, each with a characteristic power scaling law^75,76^. For instance, *α*=0.5, 0.67, and 0.75 correspond to the fluctuation of flexible partial chain, worm-like micelle, and the bending of semiflexible polymer, respectively. Notably, the high-frequency power scaling behaviour has been exploited to investigate the biophysical properties of the cytoskeletal and extracellular network of various cell conditions^42-45^. For the tissue specimens investigated here, the behaviour of reaching an elastic plateau followed by power scaling law tends to repeat at higher frequency regimes (beyond the initial Regimes I–III), where the corresponding *G*^0^, *ω*^0^, *γ*, and *α* parameters can be extracted for each repetition.

From the mechanobiological perspective, wideband micromechanical spectroscopy provides unique ‘viscoelastic fingerprints’ that characterize a wide array of tissue microenvironments. Several characteristic spectral signatures emerge from our analysis of fibrin constructs, whole blood clots, breast tissues, and bones. The first type of spectral signature is observed in acellular fibrin construct (Fig. 1d) and benign fibrous breast tissue (Fig. 3b), where the power scaling of *G*″∼*ω*^*γ*^ in Regime III follows two distinct scaling laws, with a steeper increase (i.e., larger *γ*) at higher frequency. These spectra exhibit characteristics frequency-dependent behaviours that are mostly consistent with prior studies of homogeneous fibrous polymer networks^36,39-41^. In contrast, the second type of spectral signature is observed in highly cellular tissues, including the tumour epithelium (Fig. 3h, i) and RBC-rich whole blood clots (Fig. 2). These spectra are characterized by a relatively viscous behaviour, where the viscous modulus remains close to, or even exceeds, the elastic modulus at higher frequencies. In these predominantly cellular microenvironments, the measured viscoelastic behaviour is likely dominated by the dynamics of cell surfaces, which are surrounded on either side by the cytoplasm and the interstitial fluid. The third type of spectral signature is characteristic to adipose tissue (Fig. 3e), where the delayed elastic plateau and the single steep power scaling of *G*″ result in a distinct inverted triangle shape of the spectra. This behaviour is likely governed by the viscoelastic properties of lipid, since adipose tissue is composed of densely packed adipocytes whose intracellular space is primarily occupied by lipid droplets. The last type of spectral signature is observed where fibrous stroma meets cells, including the breast tumour-stroma interface (Fig. 3d, g, i) and the cortical bone (Fig. 4b). These spectra exhibit a characteristic undulation pattern in the viscous modulus, where new elastic plateaus are repeatedly reached after ∼1 decade of power scaling of *G*″ and |*G*\*|. We hypothesize that the characteristic undulation pattern in the viscous behaviour is likely an outcome of the interaction of cell surfaces with the surrounding extracellular fibrillar network, given that this behaviour is, to the extent of our results, strictly observed where cells can directly interact with the fibrous ECM.

The wideband viscoelastic spectral signatures that emerge from various tissue microenvironments has the potential to enable novel mechano-pathological investigation and micromechanics-based clinical prognostication. In the present study, we implement a frequency regime-based analysis to extract a set of spectroscopic parameters that uniquely defines each wideband spectra. These spectroscopic parameters can provide a high-dimensional parameter space for multivariate analysis and classification, in conjunction with other known molecular biomarkers and clinical prognostic indicators. The spectral analysis of the full frequency-dependent elastic, viscous, and complex shear moduli with wideband SHEARS will offer comprehensive profiling of the tissue viscoelastic landscape for a variety of pathologies and disease conditions. The ability to access information across the broad frequency spectrum, which has thus far been inaccessible in fresh unprocessed tissue, will likely unlock new paths toward improved disease prognostication and identification of therapeutic targets.

## Methods

### Wideband SHEARS system and data acquisition

Schematic of the optical setup for the wideband SHEARS system is shown in Fig. 1a. A fibre-coupled diode laser with wavelength 637 nm (Coherent, OBIS FP 637LX) was collimated, linearly polarized, resized to a beam diameter of 1 mm, and focused by an objective lens (convex doublet, focal length 30 mm) to a spot size of 14 μm at the sample. Backscattered light was collected by the same objective lens through an open aperture of 9 mm in diameter, then, captured by a high-speed CMOS camera (Photron, Mini AX200 type 900k) through a tube lens (focal length 400 mm). Acquisition of speckle time series was accomplished with a manufacturer-provided camera control software (Photron FASTCAM Viewer 4), which enabled control of camera ROI, frame rate, and exposure time. Samples were positioned with the focal plane of the objective lens just below the sample surface. For each measurement, we used the lowest possible exposure time at which the captured speckle intensity still occupies the full available dynamic range of the camera sensor. At the maximum allowable frame rate for a given exposure time, speckle time series was acquired over a sensor ROI of 128×32 pixels for a duration of 5 s. Compared to the previous systems described in our prior work^62-65^, notable advances were implemented here to enable wideband micromechanical spectroscopy. The absolute upper and lower frequency limits of the measured viscoelastic spectra are determined by the acquisition frame rate, *F*_s_, and duration, *τ*, of the speckle time series according to 1/*τ*<*ω*<*F*_s_. Thus, extending the measurement up to the sub-MHz frequency range necessitates a high-speed camera that can capture speckle images at sub-MHz frame rate. The Photron Mini AX200 camera can support up to 540,000 frames per second (with a sensor ROI of 128×32 pixels), several orders of magnitude faster than typical CMOS cameras for microscopy applications. However, the Mini AX200 camera also comes with a much larger sensor pixel size of 20 μm×20 μm, which presents a caveat for achieving sufficient spatial sampling to capture fully developed speckles without compromising photon collection (e.g., by closing the aperture on a zoom lens). Via a combination of illumination beam size, collection aperture size, and focal lengths of objective and tube lenses, we designed the optical system to achieve a pixel-to-speckle size ratio of 3.5 pixels/speckle. Furthermore, the high-frame rate acquisition affords shorter exposure time, necessitating higher illumination power to capture the speckle intensity fluctuation with sufficient signal-to-noise ratio. The Coherent OBIS FP 637LX laser diode provides a full power of 48 mW at the sample. We utilised the full power except for measurements in whole blood (Fig. 2), where a neutral density filter was placed in the illumination arm to achieve 10 mW at the sample to avoid excessive absorption by haemoglobin.

### Reconstruction of frequency-dependent *G*\*(*ω*)

All computation was implemented in MATLAB 2022a. Firstly, ensemble-average intensity autocorrelation function, *g*_2_(*t*), was computed from the acquired speckle time series using a contrast-normalized approach described in^90^:

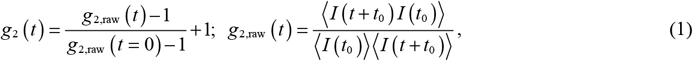

where *I* and *t* denote speckle intensity and autocorrelation time, respectively. ⟨⟩ denotes an ensemble average in space and time. Here, the ensemble includes all spatial pixels in a circular ROI concentric to the beam centre and extending 1/*e* radius of the diffused reflectance profile (DRP), and all pairs of temporal frames separated by time *t*. This ensures optimal speckle contrast in the ensemble while maximizing the number of individual speckles and temporal averaging. Then, time-dependent MSD, ⟨Δ*r*^2^(*t*)⟩, was obtained via an empirical approximation of the diffusing wave spectroscopy (DWS) formulation^62,63^:

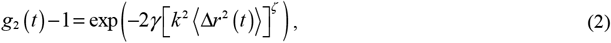

where *k* denotes wave number in the medium while *γ* and *ζ* are experimental constants that account for the optical properties of the sample. The constants *γ* and *ζ* were obtained from a lookup table derived via Monte Carlo ray tracing for a given sample optical properties, which were experimentally estimated from the radial DRP in each sample^91,92^, as previously described for laser speckle microrheology^62,63^. This approach allows SHEARS to navigate a range of unknown intrinsic optical properties in scattering biological tissues, unlike conventional DLS and DWS approaches which are limited to either the single-scattering or the multiple-scattering extremes, respectively^62,63^. Notably, a combination of *γ*=2/3 and *ζ*=1 reduces equation (2) to that of the DLS formulation while *γ*=5/3 and *ζ*=0.5 is consistent with the DWS formulation (at 180° backscattered configuration)^93,94^. In the present study, the values of *γ* and *ζ* fall between the two limits based on the optical properties of the samples summarized in Supplementary Table 2.

In principle, *G*\*(*ω*) can be obtained directly from the temporal Fourier transform of the MSD via the Generalized Stokes-Einstein Relation (GSER). In practice, since MSD is measured over a finite time domain, we used an algebraic approximation of the GSER^72-74^:

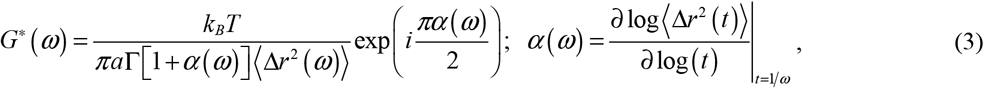

where *k*_B_, *T*, and *a* denote the Boltzmann constant, temperature, and scattering particle radius, respectively. For samples with unknown scattering particle size, *a* can be experimentally estimated from a combination of the azimuthal DRP and the relative speckle decorrelation rate of parallel versus perpendicularly polarized component of the backscattered light, as previously described^64,71^. Notably, *a* merely provides a scaling factor to the absolute values of *G*\*(*ω*) and has no effect on its frequency-dependent behaviour. The average scattering particle radii of the samples in the present study are summarized in Supplementary Table 2.

The frequency-dependent *α*(*ω*) describes the local power scaling law behaviour of the MSD, i.e., ⟨Δ*r*^2^(*t*)⟩∼*t*^*α*(*ω*)^ at *t*=1/*ω*. For optimal sampling in the frequency domain and noise performance in the log-log derivative, equation (3) was executed by first resampling the MSD linearly in the log-space *ω* domain with 30 points per decade to obtain ⟨Δ*r*^2^(*ω*)⟩, then, computing the linear regression of log⟨Δ*r*^2^(*t*)⟩ w.r.t log(*t*) over a rolling temporal window of width 7 points centred at *t*=1/*ω* to obtain *α*(*ω*). Finally, *α*(*ω*) was smoothed by a moving-average filter with a window size of 15 points before *G*\*(*ω*) was obtained via equation (3).

### Extraction of spectroscopic parameters

Spectroscopic parameters were extracted from the final reconstructed |*G*\*(*ω*)|, *G*′(*ω*), *G*″(*ω*), and *α*(*ω*) versus *ω* curves. All computation was implemented in MATLAB 2022a.

1. Transition frequency, *ω*_T_ (Regimes I and III): value of *ω* at which *α*(*ω*)=0.5.
2. Plateau frequency, *ω*^0^ (Regimes II, IV, and VI): value of *ω* at which *α*(*ω*) is at its local minima, which also corresponds to local minima of *G*″(*ω*).
3. Plateau modulus, *G*^0^ (Regimes II, IV, and VI): value of *G*′(*ω*) at *ω*=*ω*^0^.
4. Power scaling law of *G*″(*ω*), *γ* (Regimes III, V, VII): log-log slope from a linear regression of log(*G*″(*ω*)) w.r.t log(*ω*) within each regime.
5. *G*″(*ω*) power scaling transition frequency, *ω*_*γ*_ (Regime III): value of *ω* at the inflection point of the log-log derivative ∂log(*G*″(*ω*))/∂log(*ω*) within Regime III.
6. Power scaling law of |*G*\*(*ω*)|, *α* (Regimes III and V): log-log slope from a linear regression of log(|*G*\*(*ω*)|) w.r.t log(*ω*) within each regime.

### Sample preparation

Fibrin constructs (Fig. 1) were prepared with human fibrinogen plasminogen-depleted (Enzyme Research Laboratories, FIB 1) and human α-thrombin (Enzyme Research Laboratories, HT 1002a) in HBS buffer (20 mM HEPES, 135 mM NaCl, 5 mM CaCl_2_, pH 7.4) at a final concentration of 5 mg/mL fibrinogen and 2 U/mL thrombin. Fibrinogen and thrombin were separately diluted in HBS buffer to 2× the final concentrations. A volume of 150 μL of each diluted solution was added to a 96-well plate and thoroughly mixed by repeated pipetting. The plate was sealed with parafilm while the samples were allowed to polymerize at room temperature for 1 hr. Solution of unpolymerized fibrinogen (Fig. 1e) was prepared in HBS buffer at the same final concentration of 5 mg/mL fibrinogen with no thrombin. Polystyrene microspheres with diameter of 3 μm (Bangs Laboratories, PC05003) were surface functionalised with polyethylene glycol (Creative PEGWorks, mPEG-Amine, MW 5k)^95^ and added to the solution to provide scattering particles in the absence of the fibrin network structure.

Whole blood clots (Fig. 2) were prepared with patient whole blood samples from the MGH Core Laboratory (MGH IRB#2017P000419). The fibrinogen content reported by the Core Laboratory was 5.15 mg/mL for high-FIB and 2.12 mg/mL for low-FIB. Clotting was initiated by adding Kaolin (Sigma-Aldrich, K1512) and CaCl_2_ to whole blood at a final concentration of 3 μg/mL Kaolin and 14 mM CaCl_2_. The clots were prepared in a 96-well plate with a total volume of 280 μL in each well. The plate was sealed with parafilm while the samples were allowed to clot at room temperature for 1 hr.

Breast tissues (Fig. 3) were obtained from the MGH Pathology Unit following surgical tumour resection (MGH IRB#2011P000301). The specimens were stored in phosphate buffer saline at 4 °C and measured fresh within 24 hr of resection. The specimens were removed from saline, placed on top of saline-soaked gauze in a petri dish, marked with ink at four corners for subsequent co-registration with histology, and allowed to warm to room temperature. The specimens were placed on a 2-axis vernier micrometre for wideband SHEARS, where the measurement locations (arrows in Fig. 3a, c, f) were tracked w.r.t the ink marks. Measurements were taken at room temperature. Following the measurement, the specimens were fixed in 10% neutral buffered formalin, paraffin embedded, sectioned, and stained with H&E.

Bone samples (Fig. 4) were obtained from cross-sections of bovine rib and stored at 4 °C. The specimens were removed from the refrigerator, placed on top of saline-soaked gauze in a petri dish, marked with ink at four locations for subsequent co-registration with histology, and allowed to warm to room temperature. Similar to breast tissues, the wideband SHEARS measurement locations were tracked w.r.t the ink marks via vernier micrometre. Following the measurement, the specimens were fixed in 10% neutral buffered formalin, decalcified, trimmed around the ink marks, paraffin embedded, sectioned, and stained with H&E.

## Acknowledgements

The authors thank Brandon C. Matthews for technical assistance with fibrin preparations. The authors also thank Dr Jie Zhao and team at the Photopathology Core of the Wellman Center for Photomedicine for processing the tissue specimens for histology. This work was supported in part by grant R01HL142272 (PI: Nadkarni) from the National Institutes of Health.

## Author Contributions

N.L.: conception of the work; acquisition, analysis, and interpretation of all data; original draft and revision of manuscript. Z.Z.: acquisition, analysis, and interpretation of fibrin and whole blood data. Z.H.: analysis and interpretation of breast tissue data. V.B.: interpretation of breast tissue data. S.K.N.: conception of the work; acquisition, analysis, and interpretation of all data; project supervision; revision of manuscript. All authors reviewed the manuscript.

