## Supplementary Information for "Speckle rheological spectroscopy reveals wideband viscoelastic spectra of biological tissues"

### Content

Supplementary Table 1. Wideband spectroscopic parameters of fibrin constructs, whole blood clots, breast tissues, and bones

Supplementary Table 2. Optical properties and scattering particle size of fibrin constructs, whole blood clots, breast tissues, and bones

**Supplementary Table 1. Wideband spectroscopic parameters of fibrin constructs, whole blood clots, breast tissues, and bones.** Spectroscopic parameters extracted from frequency-dependent $|G^*(\omega)|$, $G'(\omega)$, $G''(\omega)$, and $\alpha(\omega)$ spectra measured by wideband SHEARS. Unless otherwise specified, values of $G$ and $\omega$ are reported in units of kPa and rad/s, respectively. Values indicate mean±standard deviation of $N$=9 (fibrin construct), 5 (whole blood clot), and 3 (breast tissue and bone) measurements.

| **Parame-ters** | **Fibrin construct** | **Whole blood clot** | | **Breast tissue** | | | | | | | **Bone** | |
|---|---|---|---|---|---|---|---|---|---|---|---|---|
| | | **High FIB** | **Low FIB** | **Fibrous** | **Adipose** | **Fibrous tumour stroma** | | | **Tumour epithelium** | | **Cortical** | **Trabecular** |
| | (Fig. 1d) | (Fig. 2a) | (Fig. 2b) | (Fig. 3b) | (Fig. 3e) | (Fig. 3d) | (Fig. 3g) | (Fig. 3i) | (Fig. 3j) | (Fig. 3h) | (Fig. 4b) | (Fig. 4c) |
| Regime I | | | | | | | | | | | | |
| $\omega_T$ | 20 ± 3 | 0.36 ± 0.09 | - | 13 ± 4 | 100 ± 30 | 30 ± 10 | 18 ± 3 | 16 ± 2 | 20 ± 10 | 8 ± 7 | 30 ± 20 | 23 ± 7 |
| Regime II | | | | | | | | | | | | |
| $\omega^0_{II}$ | 300 ± 100 | 3.1 ± 0.7 | 0.8 ± 0.2 | 210 ± 20 | 1500 ± 200 | 100 ± 20 | 86 ± 3 | 100 ± 30 | 100 ± 40 | 50 ± 20 | 1720 ± 70 | 140 ± 20 |
| $G^0_{II}$ | 0.7 ± 0.2 | 0.6 ± 0.2 Pa | 0.3 ± 0.2 Pa | 3.5 ± 0.6 | 1.27 ± 0.06 | 4.6 ± 0.9 | 0.90 ± 0.09 | 0.61 ± 0.05 | 500 ± 100 | 0.3 ± 0.2 | 5000 ± 2000 | 300 ± 100 |
| Regime III | | | | | | | | | | | | |
| $\gamma_{IIIa}$ | 0.4 ± 0.1 | 0.60 ± 0.01 | 0.68 ± 0.05 | 0.20 ± 0.03 | 0.8 ± 0.1 | 0.33 ± 0.09 | 0.44 ± 0.03 | 0.4 ± 0.1 | 0.45 ± 0.02 | 0.62 ± 0.03 | 0.56 ± 0.09 | 0.1 ± 0.1 |
| $\alpha_{IIIa}$ | - | - | - | - | 0.44 ± 0.05 | 0.22 ± 0.06 | 0.27 ± 0.05 | 0.25 ± 0.05 | 0.40 ± 0.01 | - | - | 0.16 ± 0.03 |
| $\omega_{T,III}$ | - | 110 ± 50 | 40 ± 30 | - | - | - | - | - | - | - | - | - |
| $\omega_\gamma$ | 2420 ± 70 | 160 ± 30 | 200 ± 30 | 2000 ± 200 | - | - | - | - | - | 900 ± 200 | - | - |
| $\gamma_{IIIb}$ | 1.2 ± 0.1 | 0.83 ± 0.01 | 0.80 ± 0.02 | 0.73 ± 0.05 | - | - | - | - | - | 0.51 ± 0.09 | - | - |
| $\alpha_{IIIb}$ | 0.50 ± 0.09 | 0.83 ± 0.01 | 0.80 ± 0.02 | 0.33 ± 0.05 | - | - | - | - | - | 0.51 ± 0.09 | - | - |
| Regime IV | | | | | | | | | | | | |
| $\omega^0_{IV}\times10^3$ | - | - | - | 200 ± 100 | - | 2.3 ± 0.7 | 3.1 ± 0.3 | 2 ± 1 | 3.3 ± 0.4 | - | 90 ± 30 | 1.5 ± 0.2 |
| $G^0_{IV}$ | - | - | - | 13 ± 2 | - | 8 ± 2 | 2.1 ± 0.4 | 1.2 ± 0.6 | 1.7 ± 0.4 | - | 6000 ± 2000 | 500 ± 200 |
| Regime V | | | | | | | | | | | | |
| $\gamma_V$ | - | - | - | 0.5 ± 0.1 | - | 0.55 ± 0.06 | 0.58 ± 0.06 | 0.3 ± 0.2 | 0.7 ± 0.1 | - | 0.2 ± 0.2 | 0.78 ± 0.06 |
| $\alpha_V$ | - | - | - | - | - | 0.26 ± 0.01 | 0.28 ± 0.07 | 0.26 ± 0.06 | 0.45 ± 0.07 | - | - | 0.44 ± 0.07 |
| Regime VI | | | | | | | | | | | | |
| $\omega^0_{VI}\times10^3$ | - | - | - | - | - | 130 ± 40 | 90 ± 30 | - | - | - | - | - |
| $G^0_{VI}$ | - | - | - | - | - | 18 ± 5 | 5 ± 2 | - | - | - | - | - |
| Regime VII | | | | | | | | | | | | |
| $\gamma_{VII}$ | - | - | - | - | - | 0.6 ± 0.3 | 0.35 ± 0.01 | - | - | - | - | - |

**Supplementary Table 2. Optical properties and scattering particle size of fibrin constructs, whole blood clots, breast tissues, and bones.** Estimated material parameters used in the reconstruction of frequency-dependent $G^*(\omega)$ as detailed in Methods. $\mu_s'$: reduced scattering coefficient, $\mu_a$: absorption coefficient, $a$: sphere-equivalent hydrodynamic radius.

| | **Optical properties** | **Particle radius, *a*** |
|---|---|---|
| **Fibrin construct** (Fig. 1) | $\mu_s'$ 0.6 $mm^{-1}$ | 0.18 µm |
| **Whole blood clot** (Fig. 2) | $\mu_s'$ 2.1 $mm^{-1}$, $\mu_a$ 0.1 $mm^{-1}$ | 2.88 µm$^{\dagger}$ |
| **Breast tissue** (Fig. 3) | $\mu_s'$ 1.5–3.2 $mm^{-1}$* | 0.1 µm$^{\dagger\dagger}$ |
| **Bone** (Fig. 4) | $\mu_s'$ 4.8 $mm^{-1}$ (cortical) | 12 nm$^{\dagger\dagger\dagger}$ |
| | $\mu_s'$ 3.5 $mm^{-1}$ (trabecular) | |

$^{\dagger}$Based on the hydrodynamic size of RBC[1].

$^{\dagger\dagger}$Based on prior study in breast cancer specimens[2].

$^{\dagger\dagger\dagger}$Based on the crystal size of bone minerals[3].

*The fibrous stroma tends to be more highly scattering than the cellular regions.
